# More affordable and effective noninvasive SNP genotyping using high-throughput amplicon sequencing

**DOI:** 10.1101/776492

**Authors:** Charlotte E. Eriksson, Joel Ruprecht, Taal Levi

## Abstract

Non-invasive genotyping methods have become key elements of wildlife research over the last two decades, but their widespread adoption is limited by high costs, low success rates, and high error rates. The information lost when genotyping success is low may lead to decreased precision in animal population densities which could misguide conservation and management actions. Single nucleotide polymorphisms (SNPs) provide a promising alternative to traditionally used microsatellites as SNPs allow amplification of shorter DNA fragments, are less prone to genotyping errors, and produce results that are easily shared among laboratories. Here, we outline a detailed protocol for cost-effective and accurate noninvasive SNP genotyping using highly multiplexed amplicon sequencing optimized for degraded DNA. We validated this method for individual identification by genotyping 216 scats, 18 hairs and 15 tissues from coyotes (*Canis latrans*). Our genotyping success rate for scat samples was 93%, and 100% for hair and tissue, representing a substantial increase compared to previous microsatellite-based studies at a cost of under $5 per PCR replicate (excluding labor). The accuracy of the genotypes was further corroborated in that genotypes from scats matching known, GPS-collared coyotes were always located within the territory of the known individual. We also show that different levels of multiplexing produced similar results, but that PCR product cleanup strategies can have substantial effects on genotyping success. By making noninvasive genotyping more affordable, accurate, and efficient, this research may allow for a substantial increase in the use of noninvasive methods to monitor and conserve free-ranging wildlife populations.

## Introduction

Noninvasive genetic sampling of wildlife, typically using scats or hair, is now widely used for monitoring population densities and trends for rare or elusive species (Kéry, Gardner, Stoeckle, Weber, & Royle, 2011; Waits & Paetkau, 2005; Wheat, Allen, Miller, Wilmers, & Levi, 2016). While the field data collection methods (Wasser et al. 2004) and statistical modeling (Royle, Chandler, Sollmann, & Gardner, 2014; Gardner, Royle, & Wegan, 2009; Kéry et al., 2011) needed to monitor wildlife populations have recently undergone a period of rapid advancement, noninvasive genotyping is still typically accomplished using the microsatellite and capillary electrophoresis methods developed in the 1990s. These techniques often suffer from low genotyping success and high error rates largely due to the low quantity and quality of target DNA that is intrinsic to noninvasive samples (Taberlet, Waits, & Luikart, 1999). While advances in high-throughput sequencing technologies are rapidly being developed within the biomedical, agricultural, and fisheries fields, these new technologies have struggled to gain traction in noninvasive wildlife research (McMahon, Teeling, & Höglund, 2014). Consequently, the benefits of high-throughput sequencing for noninvasive monitoring and conserving wildlife populations have largely yet to be realized.

Single Nucleotide Polymorphisms (SNPs) have emerged as a preferred alternative to microsatellites (Fabbri et al., 2012; Puckett & Eggert, 2016; Von Thaden et al., 2017). SNP genotyping allows for amplification of shorter DNA fragments, which enables higher amplification rates when using fragmented DNA from noninvasive sources (Morin, Luikart, & Wayne, 2004). Unlike microsatellite genotyping, which use size-based allele determination, SNP alleles are provided directly in the DNA sequence, which facilitates automated allele calling and results that are easily shared among laboratories and research groups.

Recent SNP genotyping approaches for noninvasive samples include the Fluidigm Dynamic Array platform (Fluidigm Corp) and the MassARRAY platform from Sequenom. The Fluidigm platform in particular has produced high genotyping success and low error rates for both fecal and hair samples (Kraus et al., 2015; Von Thaden et al., 2017). However, these techniques require specialized equipment often rendering them prohibitively expensive for wildlife research (Andrews, De Barba, Russello, & Waits, 2018; Carroll et al., 2018). A largely unexplored strategy is to amplify many SNPs together in multiplex PCR and sequence them using high-throughput sequencers. Such genotyping by amplicon sequencing (GBAS) is both cost-effective, amenable to high throughput, requires only standard laboratory equipment and no proprietary kits for library preparation, and is highly flexible with respect to the number of samples and SNPs processed. For example, GT-seq used for cost effective genotyping of animal tissue involves a highly multiplexed two-step PCR reaction where the initial PCR amplifies the targeted SNP loci and the second step adds uniquely identifiable barcodes to each sample (Campbell, Harmon, & Narum, 2015). After library preparation, the amplified PCR products are sequenced on an Illumina HiSeq sequencer and the unique barcodes are used to separate the raw sequences into unique samples using bioinformatic pipelines. Campbell, Harmon and Narum (2015) demonstrated that GBAS using the GT-seq protocol is highly cost-effective ($3.98 per sample) and produces high genotyping call rates (96.4%) for tissue samples. Natesh *et al.* (2019) provided a proof-of-concept that GBAS can work with noninvasive samples by using the GT-seq protocol to genotype 17 tiger (*Panthera tigris*) scats. However, operationalizing GBAS for noninvasive samples on a larger scale while maintaining high genotyping success rates at low cost requires additional steps and modifications to account for the inherent limitations of degraded DNA. Optimizing this method requires careful primer design to avoid non-target PCR products, additional assessment of how the size of the multiplex PCR and the number of cleanup steps influence genotyping success rates.

Multiplex PCR allows for fewer reactions and therefore increases cost and time efficiency while simultaneously reducing the amount of template DNA required. However, amplification bias and primer-dimer formations can make multiplexing challenging when using degraded DNA where primer dimers can become more abundant than target amplicons. Noninvasive genotyping approaches therefore often involve grouping primers into multiple separate smaller multiplexes (e.g. Aziz et al., 2017; Cegelski, Waits, & Anderson, 2003; Wultsch, Waits, & Kelly, 2014). In addition, purification of the PCR products to remove unincorporated primers and non-target sequences is a common step in amplicon sequencing workflows. However, this step varies in existing protocols in that some studies, including GT-seq, employed a single cleanup step after the index PCR (Fig. 1E) (Campbell et al., 2015; Lepais et al., 2019; Natesh et al., 2019) while others performed the cleanup after both PCR steps (Fig. 1B and E) (Aykanat, Lindqvist, Pritchard, & Primmer, 2016; Onda, Takahagi, Shimizu, Inoue, & Mochida, 2018; Sato et al., 2019). Performing only one cleanup step saves on cost and labor but may have downstream effects on the final results such as higher amounts of non-specific PCR products produced (Aykanat et al., 2016). For noninvasive samples, which are prone to non-target binding due to degraded DNA, a second cleanup may be particularly important to ensure genotyping success. Thus, there may be a tradeoff between genotyping success and cost and labor efficiency depending on the level of multiplexing and number of cleaning-steps used.

**Figure 1.**
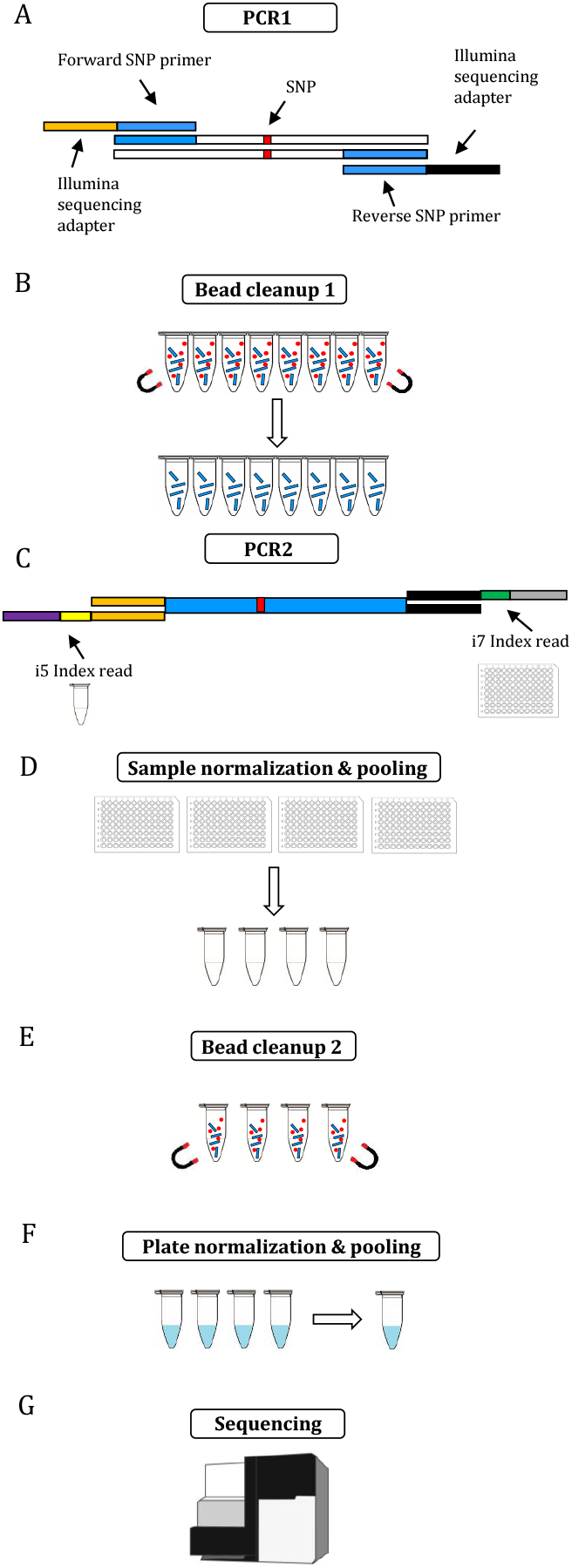
Genotyping by amplicon sequencing workflow. (A) Locus-specific primers appended with Illumina sequencing adapters are amplified in PCR1. (B) The PCR products from PCR1 are cleaned using magnetic beads to remove sequences < 120 base pairs. (C) The products from PCR1 are used as template with unique combinations of index-primers i7 (plate-specific) and i5 (sample-specific) in a second PCR reaction (PCR2). (D) The DNA concentrations are quantified for the samples in each plate and normalized to ensure even sequencing depth among samples before pooling. (E) A second round of magnetic bead-cleanup is performed to remove primer dimers and free index primers resulting from PCR2. (F) The DNA from each plate is quantified and normalized before pooling to produce even sequencing depths among the plates. (G) The pooled libraries are sent off for sequencing.

In addition, index-misassignment (hereafter, index-jumping) is a known issue of Illumina platforms (Illumina Inc, 2017). Index-jumping is thought to occur when residual free adapters or index primers are present in multiplexed sequencing libraries which can cause misassignment of indices to the wrong samples (Costello et al., 2018). For most applications, the downstream effects of this is minimal (Illumina Inc, 2017); however, for studies relying on low-coverage genomic data stemming from degraded DNA samples, even small amounts of incorrectly assigned reads could lead to false inferences (Valk, Vezzi, Ormestad, Dalén, & Guschanski, 2019). Employing strategies to minimize index jumping, such as removing free adapters by cleaning the PCR products, is therefore crucial for accurate noninvasive genotyping.

Here, we outline a detailed protocol for cost-effective and accurate noninvasive genotyping using amplicon sequencing optimized for degraded DNA. We conducted multiple experiments to refine and optimize the method in order to maximize genotyping success (Fig. 1). We used scats from known, GPS-collared coyotes (*Canis latrans)* to develop the technique and validated our results by confirming the genotypes from scats matched the genotypes from hair and tissue of the same individuals. We further demonstrate the utility of this technique for individual identification of coyotes by applying our method to 207 scat samples acquired noninvasively using detection dogs. The accuracy of the genotypes was further corroborated in that genotypes from scats matching the known GPS-collared coyotes were always located within the territory of the known individual. This optimized protocol allows for lower-cost genotypes with substantially higher success rates in order to monitor and conserve free-ranging wildlife populations.

## Methods

### Samples

We used detection dogs (Wasser et al., 2004) to locate coyote scats within and around Starkey Experimental Forest and Range in northeastern Oregon, USA. Scat samples were collected between 6-26 June 2017, georeferenced, and stored in triplicate paper bags. Within 72 hours of being collected, scats were desiccated in a drying oven for 24 hours at 40°C (Murphy et al. 2000). Additionally, coyotes were captured for a concurrent telemetry-based study using padded foothold traps monitored every 12 hours. Captured individuals were immobilized with 10 mg/kg Telazol® (tiletamine hydrochloride-zolazepam hydrochloride), given a unique ear tag, and fit with a GPS collar. We kept tissues from each ear tag punch as positive controls. The tissue samples were stored in vials with silica desiccant or 95% ethanol. Hair samples and scats (if present) were collected from captured coyotes and stored in paper envelopes. All animal handling was performed in accordance with protocols approved by the USDA Forest Service, Starkey Experimental Forest Institutional Animal Care and Use Committee (IACUC No. 92-F-0004; protocol #STKY-16-01) and followed the guidelines of the American Society of Mammalogists for the use of wild mammals in research (Sikes & Gannon, 2011).

We used a razor blade to scrape three small pieces (approximately 0.25 g in total) from the outside layer of each scat and placed them in a 2 mL Eppendorf tube. Approximately 10 hairs from each hair sample were placed in a 2 mL tube. We followed strict procedures to avoid contamination of reagents and between samples. We performed DNA extractions and PCR setup of scat and hair samples in HEPA-filtered and UV-irradiated PCR cabinets (Air Science LLC, Fort Meyers, FL) within a laboratory dedicated to processing degraded DNA (pre-PCR laboratory). We used pipettes that were only used for noninvasive samples with filter tips.

Coyote tissue samples were processed in separate extraction batches using a different PCR cabinet and pipettes to prevent contamination of the lower quality DNA samples (scats and hair). We performed all DNA extractions in small batches (11-15 samples per batch) using the DNeasy Blood and Tissue Kit (Qiagen, USA), following the manufacturer’s specifications, and included one extraction blank per batch to monitor for cross-contamination. To test for cross-species amplification of our SNP primers, we additionally extracted tissues from prey species that may be present in coyote scats. Prey species included elk (*Cervus canadensis*), deer (*Odocoileus* species), Columbian ground squirrel (*Urocitellus columbianus*), snowshoe hare (*Lepus americanus*), Deer mouse (*Peromyscus maniculatus*), montane vole (*Microtus montanus*), long-tailed vole (*Microtus longicaudus*), and bushy-tailed woodrat (*Neotoma cinerea*).

As part of a larger diet study, we used DNA metabarcoding to confirm carnivore species assignment of each scat prior to genotyping. We amplified part of the mitochondrial 12S gene using slightly modified vertebrate primers 12SV5F and 12SV5R used in Eriksson, Moriarty, Linnell, and Levi (2019), adapted from Riaz, Shehzad and Viari (2011) (see supplemental material (S1) for detailed methods). Since our 12S primers cannot distinguish between coyote and wolf (*Canis lupus),* we included SNPs with wolf-specific alleles in our panel of SNP primers (Monzón, Kays, & Dykhuizen, 2014).

### Primer Design

We designed locus-specific primers based on SNP positions and flanking regions from (Monzón, Kays, & Dykhuizen, 2014; Monzón, 2014). We used Primer3Plus (Untergasser et al., 2012) or PrimerBLAST (https://www.ncbi.nlm.nih.gov/tools/primer-blast/) to design primers optimized for multiplex success using the following parameters: 1) primer size of 19-23 bp, 2) amplicon size range of 70-120 bp, 3) melting temperature of 58-61°C with a 2°C maximum difference between the forward and reverse primer, and 4) 40-60% GC content. We avoided regions containing more than 2 di-nucleotide repeats (e.g. ATATAT) or homopolymeric regions longer than 4 bp (e.g. GGGG) due to the increased risk of PCR error. We checked for secondary structures using OligoAnalyzer 3.1 (Integrated DNA Technologies) and accepted only primers with ΔG values that were more positive than −7 kcal/mole for any self-dimers or heterodimers.

To avoid potential hairpins we selected primers with ΔG values larger than −2 kcal/mole and with temperatures that would not exist during PCR setup or in the thermocycler. We were particularly careful to avoid secondary structures (or complementarity among primers) at or near the 3’ end of the primers. We evaluated primer specificity in-silico by searching for the primer sequences in the dog genome (CanFam3.1) using BLAT (http://genome.ucsc.edu/) and BLAST (www.ncbi.nlm.nih.gov/blast) to check for complementarity of potential prey DNA present in the scat. Finally, we used the Multiple Primer Analyzer (thermofisherscientific.com) to check for potential formation of secondary structures among different primers pairs. We redesigned primers if potential secondary formations had ΔG values of less than −10 kcal/mole. Each primer was appended with overhang adapter sequences (33-34 bp) which are compatible with the index barcodes and Illumina sequencing adapters. The primers were ordered from Integrated DNA Technologies (IDT) in 100 μM concentrations using standard desalting purification. All primer pairs were tested in singleplex PCRs and visualized on a 1% agarose gel to ensure amplification, correct product size, and to optimize annealing temperatures and initial primer concentrations.

Illumina sequencing adapters P7 and P5 containing unique 8 bp i7 and i5 barcodes (hereafter, index primers) were ordered in plate format from IDT in 100 μM concentrations using standard desalting purification. We prepared multiple 10 μM stock solutions of the index primers in 8-well (i5 indices) and 12-well tube strips (i7 indices). For this study we used 16 unique i5 primers and 24 unique i7 primers creating 384 unique pairwise barcode combinations. We therefore refer to a sequencing library as 4 96-well plates (96 x 4 = 384). A larger number of samples is possible by ordering more unique index primers.

### Amplicon sequencing – Multiplex test

The first PCR consisted of 2 μL (0.2-0.3 μM final concentration) of each locus-specific primer (n=26), 10 μL Amplitaq Gold Master mix (Applied Biosystems, Foster City, CA), 7 μL water and 1 μL DNA template for a total volume of 20 μL. After an initial denaturation for 10 min at 95°C, the PCR cycling conditions were [95°C for 30 s, 56 °C for 30 s, 72°C for 30 s] for 45 cycles (35 cycles for tissue samples) followed by a 7 min final extension at 72°C. We performed triplicates of each PCR reaction and included three no template controls (NTCs) per 96-well plate to monitor for contamination.

We conducted a test to determine if multiplexing fewer primers per PCR would increase our genotyping success rate. We compared three multiplex treatments: all 26 loci (M1) in one multiplex, two multiplexes targeting 12 and 14 loci in each (M2), and three multiplexes with 8, 8 and 9 loci in each (M3). Following PCR1, the M2 and M3 groups were pooled into single 96 well-plates. We performed bead cleanup of each PCR plate using PCRClean DX (Aline Biosciences, USA) in 1.8x reaction volume to remove any sequences shorter than 120 bp (primer dimers, unincorporated dNTPs).

In the second PCR, 11 μL of the purified products from PCR1 were used as template in 27.5 μL reactions containing a unique combination of 2 μL (0.7 μM final concentration) each of P7 and P5 index primers, and 12.5 μL Q5 DNA Polymerase (New England BioLabs, Ipswich, MA). The index primers and polymerase were added to the plates in the pre-PCR laboratory and then brought to the post-PCR room where the template was added. After initial denaturation of 98°C for 30 s, the cycling conditions were [98°C for 10 s, 55°C for 30 s, 65°C for 30 s] for 8 cycles followed by a 5 min final extension at 65°C.

To ensure even sequencing depth across samples, we quantified DNA concentration using a fluorescence microplate reader with the AccuBlue dsDNA Quantitation Kit (Biotium, Hayward, CA) and normalized each sample accordingly. Following normalization, 3 μL from each sample per 96-well plate were pooled into a 0.65 mL Eppendorf tube. We then performed a second round of bead-cleanup, quantified each tube using a Qubit 2.0 fluorometer (Life Technologies, Carlsbad, CA), normalized and pooled the tubes before sending the final library pool for 150bp single-end sequencing on an Illumina HiSeq 3000 at the Center for Genome Research and Biocomputing, Oregon State University. Library size distribution was checked prior to sequencing using a High Sensitivity D5000 DNA ScreenTape assay on an Agilent Tapestation 4200 (Agilent Technologies, Santa Clara, CA).

### Amplicon sequencing – Cleanup experiment

We experimentally evaluated whether the more expensive and time-consuming double cleanup step was necessary by testing for differences in genotyping success rates and the average proportion of mis-assigned reads. We varied the cleanup workflow with two treatments: (1) bead-cleanup of each PCR product individually after PCR1 and again after the final pooling (hereafter, 2 bead-cleanups) (Fig. 1B and E), and (2) only cleaning after pooling (hereafter, 1 bead-cleanup (Fig. 1E). To assess the effects on genotyping success we prepared 23 coyote scats (two replicates each) with two cleanups and repeated the same samples but with only one cleanup. This allowed us to control for sample quality while assessing the effects of the two cleanup treatments.

The occurrence of index-jumping is more challenging to detect when amplifying similar samples such as a single species with the same primer pool added to each sample. To determine the rate of mis-assiened reads we therefore added samples and primers from two additional species, cougar (*Puma concolor*) and black bear (*Ursus americanus*) so that index jumping would be immediately apparent and quantifiable based on the rate of which cougar and black bear sequences appeared in coyote samples. These experiments were conducted using two additional libraries with all PCR reactions performed in triplicate with three NTCs per plate unless otherwise stated. The first library used two bead cleanups with 99 coyote samples (with 26 SNP primers) and 25 cougar tissues (with 25 SNP primers) (Library A). The second library consisted of 62 coyote scats prepared with two bead-cleanups, 23 coyote scats that were prepared following both one-cleanup and two-cleanup workflows (the same samples as used to determine the effects of cleanup on genotyping success described above), and 31 black bear samples (with 26 SNP primers) prepared with one cleanup (Library B). This allowed us to test whether coyote samples cleaned only once would be more infiltrated by bear samples cleaned only once. Libraries A and B were sequenced on two separate sequencing runs. We assessed the rate at which cougar and bear sequences appeared in coyote samples to quantify the effect of cleanup treatment on the average proportion of mis-assigned reads within and among plates.

### Data Analysis

The raw sequencing reads were demultiplexed and split into individual samples based on their unique index combinations using Illumina software (bcl2fastq). We used perl scripts by Campbell, Harmon and Narum (2015) to first assign a locus to each read based on the forward primer sequence, then matching the read to a specified in-silico probe targeting a region immediately surrounding the SNP (6 bp in either direction). Only sequences including both the forward primer and the in-silico probe (hereafter on-target reads) contributed to producing genotypes. Reads containing either allele were then counted and the ratio of allele 1 to allele 2 was used to assign a genotype using the GTseq_Genotyper_v3.pl PERL script by Campbell et al. (2015). Briefly, a ratio of: >10 was called homozygous for allele 1, <0.1 homozygous for allele 2, and < 2 were called heterozygous.

We constructed consensus genotypes for each sample based on the three replicates. We required a heterozygous genotype to be seen in at least two replicates and all three replicates for a homozygous genotype. We screened the replicates for loci with low read abundance and removed genotypes that were close to the NTCs. We quantified genotyping error rates (allelic dropout and false alleles) based on mismatches among the sample replicates and by comparing genotypes produced in noninvasive samples to reference genotypes from tissue samples according to Broquet and Petit (2004). We discarded samples with > 20% loci missing or with an allelic dropout rate of > 0.4. We estimated the probability of identity (PID) and probability of identity among siblings (PIDsibs) using GenAlEx v 6.5 (Peakall & Smouse, 2012). We used the R package allelematch to identify unique individuals (Galpern, Manseau, Hettinga, Smith, & Wilson, 2012).

## Results

### Genotyping success

We genotyped 207 randomly selected scat samples collected during the detection dog surveys in addition to 18 hair, 15 tissue, and 9 scat samples from the captured coyotes. Out of 216 scat samples, we discarded 13 (all from the detection dog surveys) due to missing data (missing > 5 loci), and 2 with allelic dropout rates of > 0.4, leaving 201 samples resulting in a genotyping success rate of 93%. Call rates for the consensus genotypes from the scat samples were high with 90% of loci genotyped in 198 samples (91% of samples) and all loci yielding genotypes in 169 samples (78% of samples) (Fig. 2). The genotyping success rate for hair and tissue was 100%.

**Figure 2.**
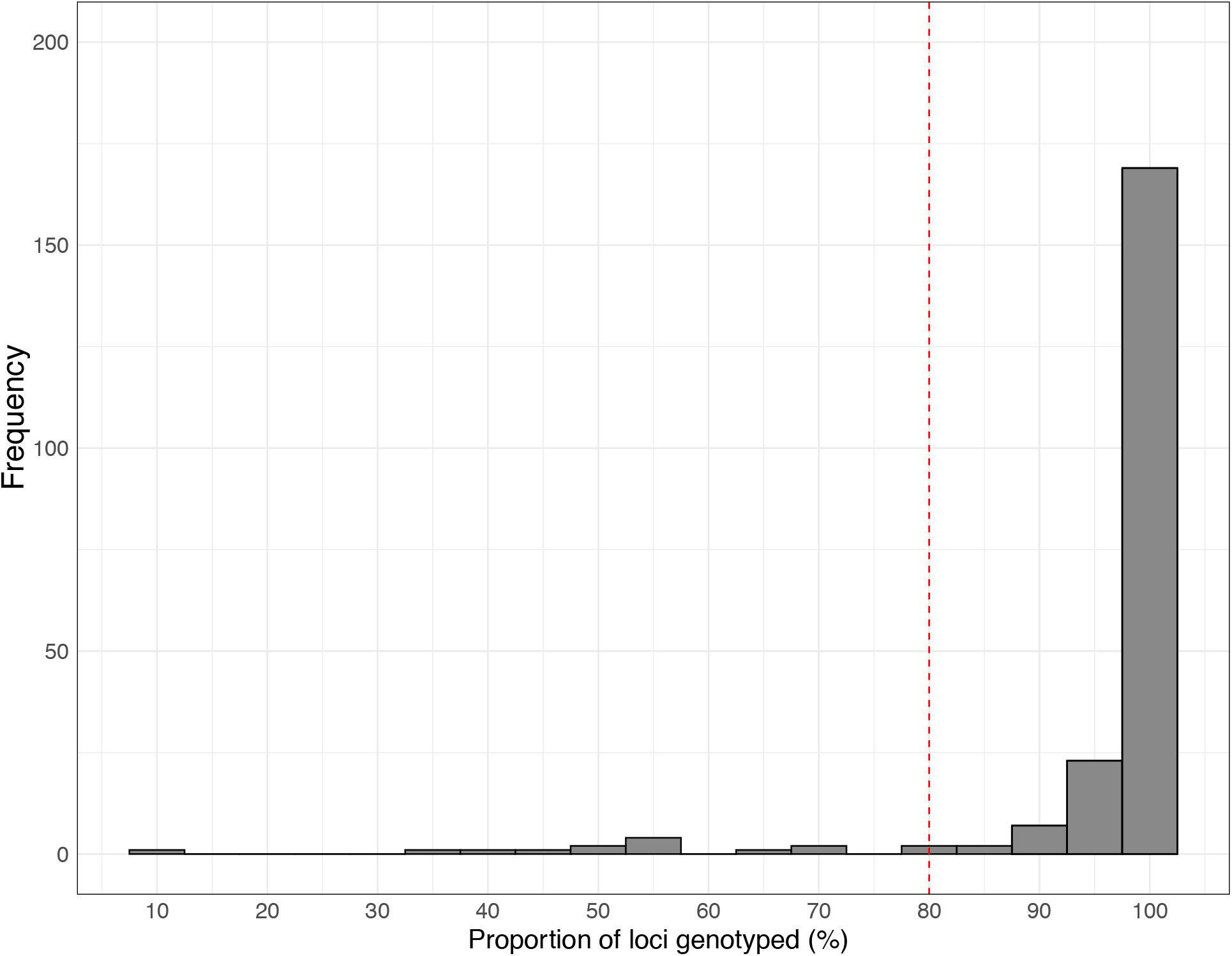
Histogram showing the genotype call rates for consensus genotypes of coyote (*Canis latrans)* scat samples. The red line represents the filtering threshold of proportion of loci required to retain a sample. Samples that produced a consensus genotype in less than 80% of loci were discarded.

There were no disagreements between genotypes from the hair, tissue and scat samples from the captured coyotes. Error rates based on comparing noninvasive samples (hair and/or scat) from known individuals to reference samples (tissue) were therefore 0%. The false allele and allelic dropout rates calculated from scat sample replicates were 0.001 and 0.01 respectively. No errors were found in the hair replicates. The probability of identity for all loci combined was 0.000001 and 0.001 among siblings. None of the tested prey species contained both the forward primer and in-silico probe required to produce a genotype. All genotypes produced were therefore from coyotes free of contamination from prey species.

### Individual identification

Three of the putative coyote scats were wolf scats and were therefore removed. A total of 90 individual coyotes were found from the remaining 198 scats. Eleven of the 17 collared coyotes were detected from scats collected during the detection dog surveys (Fig. 3). The location of the matching scat always corresponded to the known territory of the individual coyote based on 95% kernel density estimates (Fig. 3).

**Figure 3.**
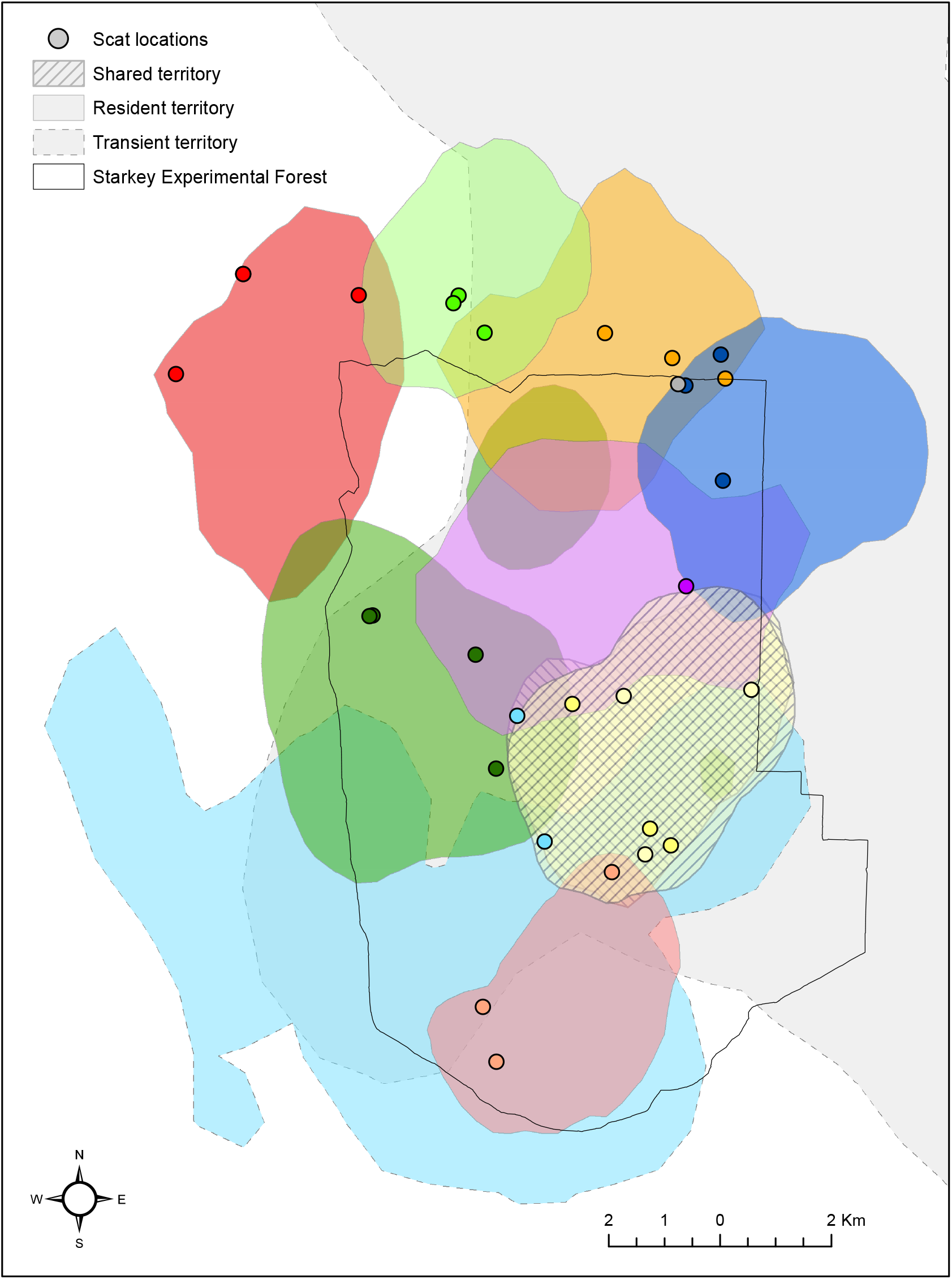
Study area with territories of GPS-collared coyotes (*Canis latrans)* and the matching scat samples collected by scat detection dogs. A similarly colored scat sample and underlying territory indicates a matching genotype.

### Multiplex test

The average proportion of on-target reads varied slightly among the three multiplex treatments with 78.3%, 85.9 % and 83.6% respectively for M1 (all 26 loci), M2 (12 and 14 loci) and M3 (8, 8, and 9 loci). Genotyping success rates were therefore also similar (81.7%, 83.9%, 81.7% respectively) irrespective of the number of primers multiplexed.

### Cleanup strategies and index jumping

The two bead-cleanup strategies had substantial effects on genotyping success. The samples with two cleanups produced mean DNA concentrations of 19.8 ng/uL (± 6.6 SD) compared to 39.7 ng/uL (± 6.8 SD) for the identical samples cleaned only once. This resulted in multiple negative downstream effects with the samples cleaned once producing on average two-thirds fewer on-target reads than the same samples cleaned twice (102805.2 ± 6530.1 SD vs. 320372.8 ± 23718.9 SD) (Fig. 4). In addition, the samples that were cleaned once had on average 5.26 ± 0.32 SD loci that failed to produce a genotype compared to 0.89 ± 0.26 SD for the samples cleaned twice.

**Figure 4.**
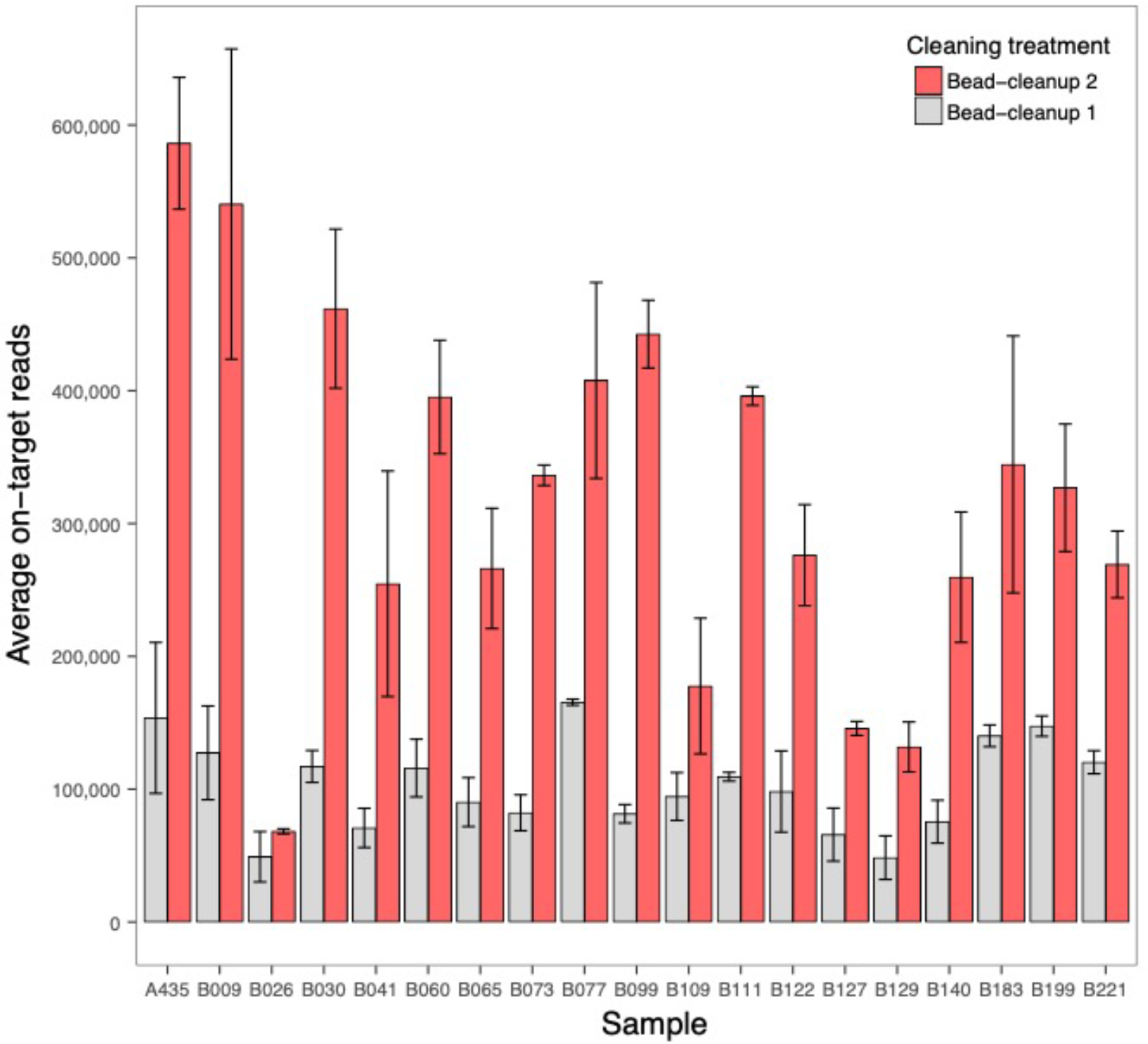
Average on-target reads (reads containing both the forward primer and in-silico probe) per scat sample and cleaning treatment. 2 Bead-cleanups involved a two-step cleaning process, first of individual samples after PCR1 and also after the samples were pooled. 1 Bead-cleanup involved only cleaning after the samples were pooled.

In addition to the effects on genotyping success, the cleanup strategies had a considerable influence on the amount of index-jumping among plates. Index-jumping was miniscule in Library A where all four plates were cleaned twice (Fig. 5A). The number of cougar reads incorrectly assigned to coyote samples ranged from 0.00037% to 0.59% of the total amount of on-target reads per sample averaged across plates.

The coyote samples that were cleaned twice in Library B had on average 0.16% - 1.64% bear reads incorrectly assigned. In comparison, the coyote samples prepared with one bead-cleanup had on average 8.72% incorrectly assigned bear reads per sample (Fig 5B). The bear samples also cleaned once had on average 6.60% incorrectly assigned coyote reads per sample. Index-jumping was higher between plates that shared either the i7 or i5 sets of index primers compared to plates that shared no indices (Fig. 5A and B).

**Figure 5.**
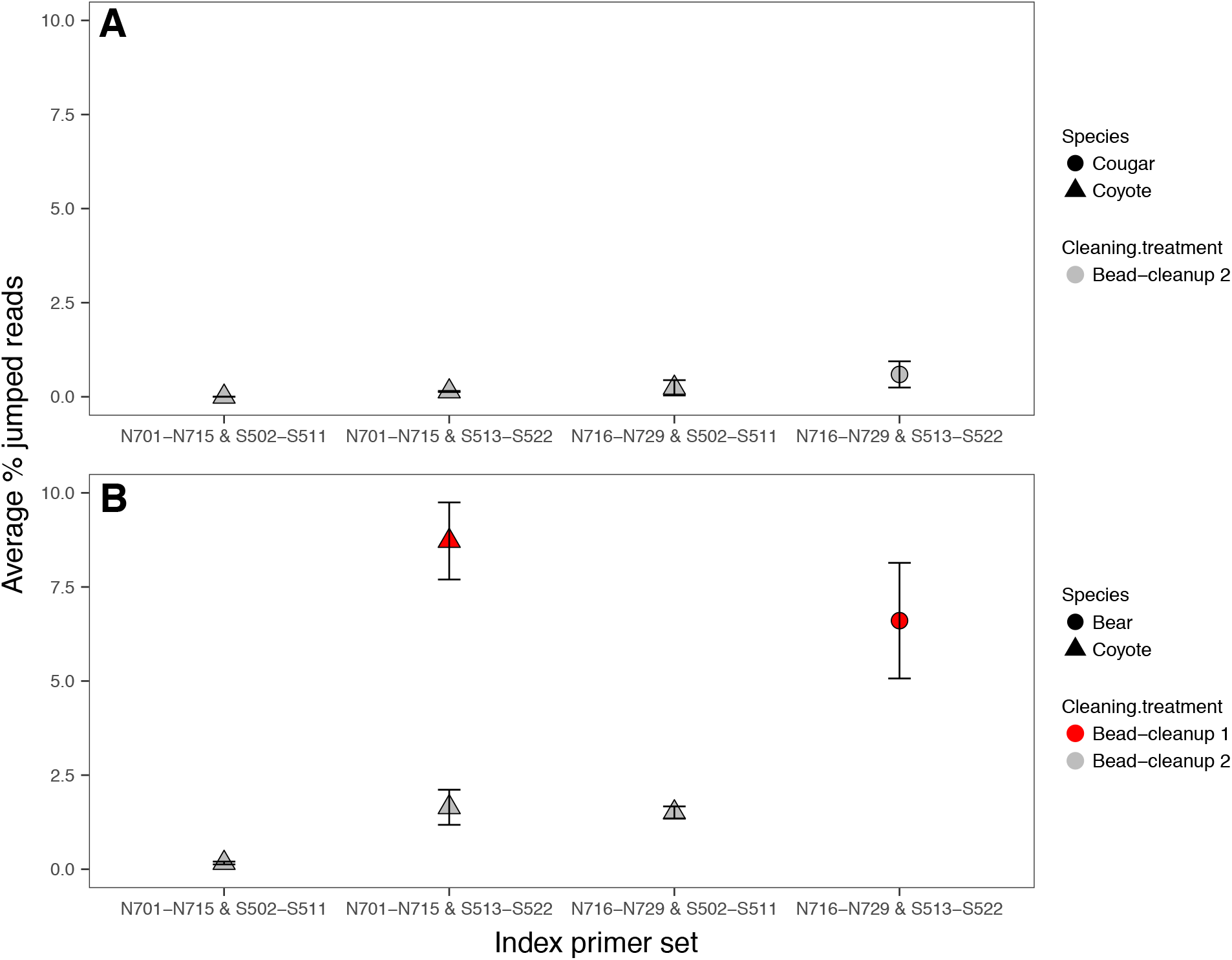
Effects of cleaning treatment on the proportion of miss-assigned reads per plate. 2 bead-cleanups involved a two-step cleaning process, first of individual samples after PCR1 and after the samples were pooled. 1 bead-cleanup involved only cleaning after the samples were pooled. A) Proportion of jumped reads within a sequencing library where all samples were cleaned twice. B) Proportion of jumped reads within a library where 62 coyote samples were cleaned twice, 23 coyote scats were prepared following both one-cleanup and two-cleanup workflows, and 31 black bear samples prepared with one cleanup.

## Discussion

Noninvasive genotyping methods have become key elements of wildlife research over the last two decades, but their widespread adoption is limited by high costs, low success rates, and high error rates. Newer SNP genotyping platforms can improve genotyping success rates but require specialized equipment and can be prohibitively expensive. High-throughput sequencing technologies are rapidly progressing and decreasing in cost. Wildlife conservationists seeking to employ these advances need to carefully adapt protocols to account for the imperfect circumstances of field biology, wild study subjects, and degraded DNA. Here, we use laboratory protocol experiments to present a cost-effective and accurate genotyping method using amplicon sequencing optimized for low-quality DNA samples.

Our genotyping success rate of 93% was considerably higher than previously published microsatellite-based studies on coyotes using scats (27.5%-65.1%) (Table S2). Our success rate is also higher than that 77% rate found by Monzón et al. (2014), which as far as we know is the only prior study to have genotyped coyote scats using SNPs. Our genotyping success rate was also higher than other recent SNP-based noninvasive genotyping studies with scats from other species using the MassArray system (59.8%) (Fitak et al. 2015), and Fluidigm platform 87% (Von Thaden *et al.,* 2017), 75% (Ekblom et al., 2018), 90% (Kraus et al., 2015). The hair samples used in this study were collected as references from captured coyotes and could therefore not be considered truly noninvasive, however, the high success rate (100%) is promising for future applications using hair-snares. Compared to the error rates of microsatellite genotyping (0-48%) (Broquet & Petit, 2004), previous research has found that SNP-based methods substantially reduce genotyping error rates to 1% (Kraus et al., 2015), 0.00038% (Norman & Spong, 2015) and 0.0029% (Bourgeois et al., 2019) when pre-selecting samples based on DNA concentration. Our total error rate for scats was 0.01% without any pre-selection based on sample quality beyond selecting randomly from the subset of scats that yielded species ID via DNA metabarcoding (85% of scats yielded a species ID).

Our overall high success rate is likely due to several factors; short target amplicons, careful primer design to minimize interactions and non-target amplification, and the high sensitivity of next-generation sequencing technologies. This high success rate coupled with the affordability of highly-multiplexed amplicon sequencing produces a substantial cost saving with further cost reductions as the price of sequencing continue to decline. The cost of generating these data from DNA extraction to sequencing are $4.87 per replicate (excluding labor) (Table S1) with the single greatest cost per sample attributed to DNA extraction ($2.0 per sample). Using a more economical extraction kit could reduce the per-sample cost further. Because we observed no disagreement among replicates for samples with 100% call rates, costs could be reduced further by reducing the number of replicate PCRs as recommended by Von Thaden et al. (2017). We did not include labor cost as these vary across institutions or primer development costs as these depend on the genomic information available for the focal species and are not a recurring cost. A further cost-reducing benefit of amplicon sequencing is the complete flexibility in the number of loci and samples genotyped. In comparison, the Fluidigm system is restricted to 96 samples genotyped at 96 loci, 48 samples at 48 loci or 192 samples at 24 loci (www.fluidigm.com). To allow for a direct cost comparison, if we genotyped 96 samples using 96 SNPs, the total cost using our amplicon sequencing approach would be $255.5, compared to $1,418 using the Fluidigm platform, and $1,560 for the MassArray system based on the cost calculations in Carroll *et al.* (2018). Our cost calculations are based on using the full capacity of a Hiseq lane; approximately 2000 samples (Campbell et al., 2015). For comparison, if only 384 samples were genotyped, the total cost would be $7.24 per sample, but sequencing smaller numbers of samples is possible by combining multiple projects onto a single sequencing lane.

The Fluidigm and MassArray platforms, as well as traditional microsatellite genotyping, do not involve any sequencing but rather produce allele calls at specific loci. This can be an advantage since no bioinformatic skills are needed, however, these technologies still rely on the design of locus-specific SNP primers. One benefit of genotyping by amplicon sequencing is that considerably more information is obtained than just the specific alleles. Depending on the length of the amplicon, approximately 50 bp on either side of the target SNP is generated. This enables highly effective primer optimization as problematic primers producing dimers, non-target binding, low or too high read abundances can be easily identified, redesigned or removed. If the primers were designed based on genomic information from a closely related species, as often is the case for non-model species, obtaining the flanking regions also allows for the discovery of additional SNPs apart from the target variant. Targeting amplicons with multiple SNPs can be used as multi-allelic loci, increasing the power of each locus for parentage assignment (Baetscher, Clemento, Ng, Anderson, & Garza, 2017).

We demonstrated that cutting costs by reducing the number of bead cleanup purifications is not worthwhile because purification of PCR products can have a considerable impact on genotyping success. Samples treated with two rounds of bead cleanup (first after PCR1 and second time after pooling) produced more on-target reads and therefore higher genotyping success rates. This is likely due to the removal of primer dimers and other non-target sequences present after PCR1. If left uncleaned, these non-target sequences also falsely inflate the DNA concentrations per sample leading to incorrect dilution of samples during the normalization step (Fig1. D). Reducing primer concentration can be another strategy to minimize primer dimers.

However, DNA concentrations from noninvasively collected samples are highly variable and optimizing primer concentrations to match the sample quality is therefore difficult. When using degraded DNA samples, non-target sequences are inevitably produced as very little target DNA is available. The additional cleaning step costs $0.30 per sample and takes approximately 1 hour per 96-well plate. The addition of a second cleanup step was also crucial in the reduction of index-jumping. By adding a different species to sequencing libraries we were able to quantify the amount of index-jumping among samples. We show that by cleaning the samples twice, index-jumping is mitigated and kept to minimal levels. Another strategy would be to use all unique i7 and i5 index primers which would reduce the risk of index-jumping even further but would increase cost.

Many facets of wildlife ecology require estimates of population density. For example, a basic requisite for conserving wildlife populations is being able to monitor trends in abundance over time, and many fundamental ecological questions related to species interactions, density-dependence, and factors affecting population growth rates require precise estimates of animal abundances. Although noninvasive genotyping may be the only practical method to estimate the densities of many species, high costs and low genotyping success rates limit its accessibility.

SNP genotyping can increase success rates, but the widespread adoption of SNPs in wildlife research has lagged because of high costs and a lack of available markers. However, recent progress in high-throughput sequencing technologies has resulted in both a rapid increase of genomic information that simplifies SNP discovery, and has reduced costs by allowing for thousands of multiplexed amplicon samples to be affordably sequenced on a single lane. Further, the higher genotyping success rates achieved here increase precision in density estimation for improved conservation and management decisions at lower cost. Our approach therefore represents a valuable contribution to address the need for an accurate, cost- and time-effective method of noninvasive genetic monitoring of free-ranging wildlife species.

## Acknowledgements

We thank Oregon Department of Fish and Wildlife for funding this work and for providing logistical support, Jennifer Allen for assistance in the lab and Matthew Peterson for bioinformatics help.

## Data accessibility

All genotyping data is available at (link).

## Author contributions

CE and TL conceived the research and designed the experiments. CE performed primer design, laboratory work and bioinformatic analysis. JR collected all the samples and produced coyote territory estimates. CE wrote the first version of the paper, all authors provided edits and comments.

## Supporting Information

Table S1 Genotyping cost

Table S2: Genotyping success rates of other studies using coyote scats

